# Predicting Enhancer-Promoter Interaction from Genomic Sequence with Deep Neural Networks

**DOI:** 10.1101/085241

**Authors:** Shashank Singh, Yang Yang, Barnabás Póczos, Jian Ma

## Abstract

In the human genome, distal enhancers are involved in regulating target genes through proximal promoters by forming enhancer-promoter interactions. Although recently developed high-throughput experimental approaches have allowed us to recognize potential enhancer-promoter interactions genome-wide, it is still largely unclear to what extent the sequence-level information encoded in our genome help guide such interactions. Here we report a new computational method (named “SPEID”) using deep learning models to predict enhancer-promoter interactions based on sequence-based features only, when the locations of putative enhancers and promoters in a particular cell type are given. Our results across six different cell types demonstrate that SPEID is effective in predicting enhancer-promoter interactions as compared to state-of-the-art methods that only use information from a single cell type. As a proof-of-principle, we also applied SPEID to identify somatic non-coding mutations in melanoma samples that may have reduced enhancer-promoter interactions in tumor genomes. This work demonstrates that deep learning models can help reveal that sequence-based features alone are sufficient to reliably predict enhancer-promoter interactions genome-wide.

## INTRODUCTION

Our understanding of how the human genome regulates complex cellular functions in a living organism is still limited. A critical challenge is to fundamentally decode the instructions encoded in the genome sequence that regulate genome organization and function. One particular aspect that we still know little about is the three-dimensional higher-order organization of the human genome in cell nucleus. The chromosomes in each human cell are folded and packaged into a nucleus with about 5µm diameter. Intriguingly, this packaging is highly organized and tightly controlled [1]. Any disruption and perturbation of the organization may lead to disease. Recent development of new high-throughput whole-genome mapping approaches such as Hi-C [2] and ChIA-PET [3, 4] has allowed us to identify genome-wide chromatin organization and interactions comprehensively. We now know that the global genome organization is more complex than previously thought, in particular, in regards to enhancer-promoter interactions. Distal regulatory enhancer elements can interact with proximal promoter regions to regulate the target gene’s expression, and mutations that change such interactions will cause target gene to be dysregulated [5–7]. In mammalian and vertebrate genomes, the promoter regions of the gene and their distal enhancers may be millions of base-pairs away from each other; and a promoter may not interact with its closest enhancer. Indeed, the mappings of global chromatin interaction based on Hi-C and ChIA-PET have shown that a significant proportion of enhancer elements skip nearby genes and interact with promoters further away in the genome by forming long-range chromatin loops [8, 9]. However, the principles encoded at the genomic sequence level underlying such organization and chromatin interaction are poorly understood.

In this work, we focus on determining whether the sequence features encoded in the genome within enhancer elements and promoter elements are sufficient to predict enhancer-promoter interactions (EPI). Although certain sequence features (e.g., CTCF binding motifs [10]) are known to be involved in mediating chromatin loops, it remains largely under-explored whether and what information encoded in the genome sequence contains important instructions for forming EPI. There exists some recent work on predicting EPI based on functional genomic features [11, 12]. In [11], a method called RIPPLE was developed using a combination of random forests and group LASSO in a multi-task learning framework to predict EPIs in multiple cell lines, using DNase-seq, histone marks, transcription factor (TF) ChIP-seq, and RNA-seq data as input features. In [12] the authors developed TargetFinder based on boosted trees to predict EPI using DNase-seq, DNA methylation, TF ChIP-seq, histone marks, CAGE, and gene expression data. From RIPPLE and TargetFinder, it is clear that signals from functional genomic data are informative to computationally distinguish EPIs from non-interacting enhancer-promoter pairs. There are also recent works that utilize functional genomic data from multiple datasets to identify EPIs [13, 14]. These studies suggest that important proteins and chemical modifications that may be involved in mediating chromatin loops for EPIs can be recognized. However, it remains unclear whether the information in genome sequences within enhancers and promoters alone is sufficient to distinguish EPIs. Indeed, no other algorithm currently exists to predict EPI using sequence-level signatures only except our own recent work called PEP [15] (which we will directly compare in this work). PEP uses a machine learning model Gradient Tree Boosting [16] based only on features from the DNA sequences of the enhancer and promoter regions. Specifically, it considers two variants, PEP-Motif, which only uses motif enrichment features for known TF binding motifs, and PEP-Word, which uses word embeddings, a recent innovation from natural language processing that allows representing (arbitrary-length) sentences from a discrete vocabulary as fixed-length numerical vectors, while retaining semantic meaning. PEP’s results show that it is possible to achieve comparable results using sequence-based features only to predict EPIs. However, it is unclear whether different models can be developed to have even better performance.

In this paper, we want to answer the following question: *if we are only given the locations of putative enhancers and promoters in a particular cell type, can we train a predictive model using deep neural networks to identify EPIs directly from the genomic sequences without using other functional genomic signals?* In the past two years, there have been many deep learning applications to regulatory genomics [17–26]. The deep learning framework has the advantage of automatically extracting useful features from the genome sequence and can capture non-linear dependencies in the sequence to predict specific functional annotations [27]. However, three-dimensional genome organization and high-order chromatin interaction of functional elements remain an unexplored area for deep learning models.

To approach this, we develop, to the best of our knowledge, the first deep learning architecture for predicting EPIs using only sequence-based features, which in turn demonstrates that the principles of regulating EPI may be largely encoded in the genome sequences within enhancer and promoter elements. We call our model SPEID (Sequence-based Promoter-Enhancer Interaction with Deep learning; pronounced “speed”). Given the location of putative enhancers and promoters (that are largely cell-type specific) in a particular cell type, SPEID can effectively predict EPI in that cell type using sequence-based features extracted from the given enhancers and promoters based on a predictive model trained for that cell type. In six different cell lines, we show that SPEID achieved better results in both AUROC and AUPR as compared to PEP and TargetFinder. We also present two approaches for using SPEID to identify sequence features that are informative for predicting EPI. While feature importance as measured by TargetFinder and PEP, which incorporate hand-crafted features, can depend on these handcrafted features, SPEID allows a more objective approach to feature identification. In addition, we demonstrate that SPEID can help identify possible important non-coding mutations that may reduce or disrupt chromatin loops in cancer genomes. We believe that SPEID has the potential to become a generic model to allow us to better understand sequence level mechanistic instructions encoded in our genome that determine long-range gene regulation in different cell types. The source code of SPEID is available at: https://github.com/ma-compbio/SPEID

## RESULTS

### Overview of the SPEID model and data

Like all deep learning models, SPEID learns a sequence of increasingly complex feature representations. Specifically, as illustrated in Fig. 1, SPEID consists of three main layers: a convolutional layer, a recurrent layer, and a dense layer. The convolution layer learns a large array of independent ‘kernels’. Kernels are short (40bp) weighted sequence patterns that are convolved with the input sequence to compute the match of that pattern at each position of the input. Hence, the convolution layer outputs, for each kernel, the match at each position of the input. The recurrent layer re-weights each kernel match, so as to learn predictive combinations of kernel features. It does this by iterating across the length of the input sequence (in both directions, in parallel), and selectively down-weighting kernel matches based on the match strength and the weights of previously observed kernel matches. Finally, the dense layer is a simple, essentially linear, classifier learned on top of the combinations of sequence features output by the recurrent layer. We assume that important sequence features may differ between enhancers and promoters, and that interactions between enhancer and promoter sequence features determine EPI. Hence, convolution layers are separate for enhancers and promoters, and the outputs of the convolution layers are concatenated before feeding into the recurrent layer.

**Figure 1:**
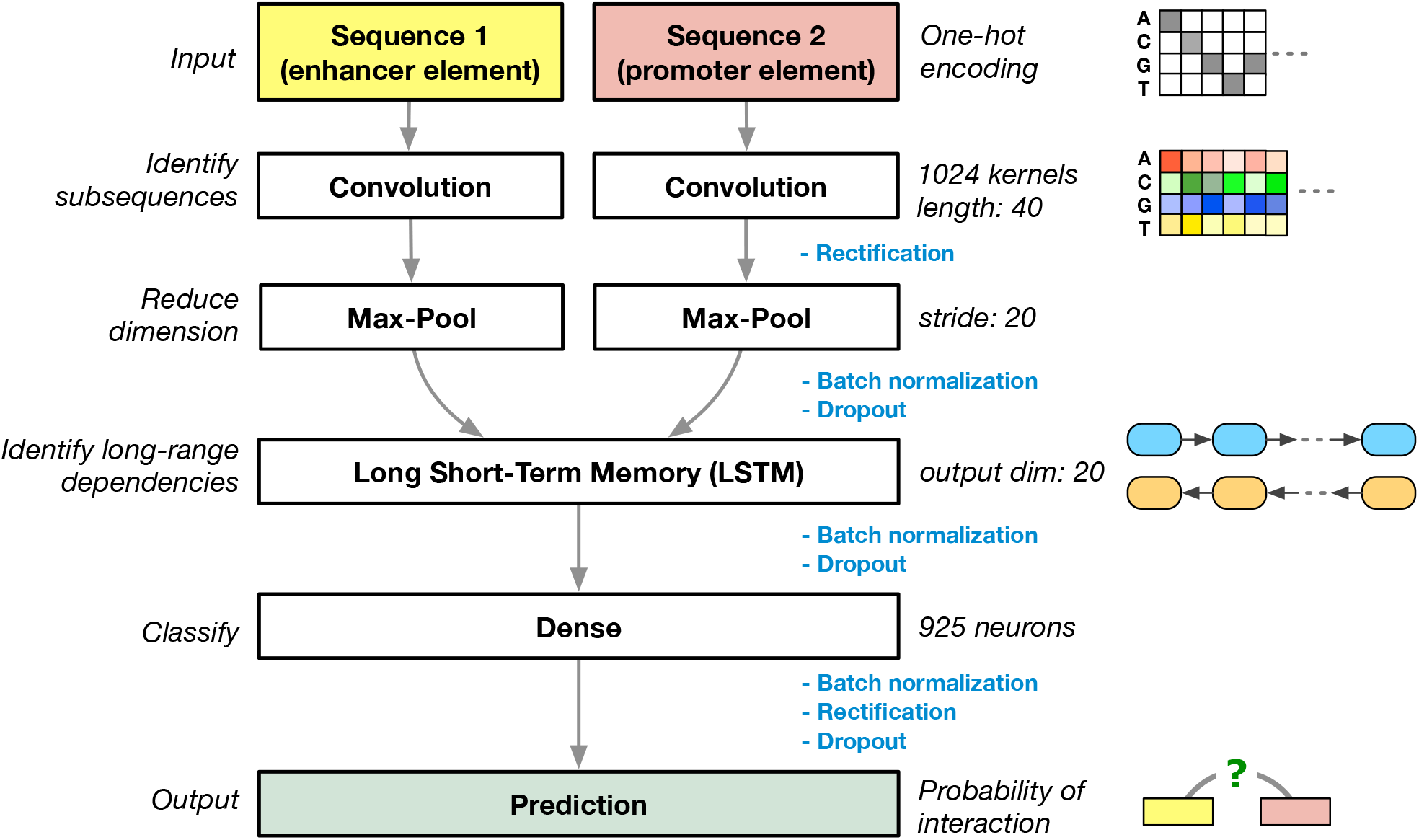
Diagram of our deep learning model SPEID to predict enhancer-promoter interactions based on sequences only. Key steps involving rectification, batch normalization, and dropout are annotated. Note that the final output step is essentially a logistic regression in SPEID which provides a probability to indicate whether the input enhancer element and promoter element would interact.

We utilized the EPI datasets previously used in TargetFinder [12], which was also used in PEP [15], for our model training and evaluation so that we can also directly compare with TargetFinder and PEP. The data include six cell lines (GM12878, HeLa-S3, HUVEC, IMR90, K562, and NHEK). Cell-line specific active enhancers and promoters were identified using annotations from the ENCODE Project [28] and Roadmap Epigenomics Project [29]. The locations of these putative enhancers and promoters are the input for SPEID for each cell line. The data for each cell line consist of enhancer-promoter pairs that are annotated as positive (interacting) or negative (non-interacting) using high-resolution genome-wide measurements of chromatin contacts in each cell line based on Hi-C [10], as used in [12]. 20 negative pairs were sampled per positive pair, under constraints on the genomic distance between the paired enhancer and promoter as described in [12], such that positive and negative pairs had similar enhancer-promoter distance distributions.

To address the problem of class imbalance, we applied data augmentation to the positive pairs (see below). The original annotated enhancers in the datasets of each cell type are mostly only a few hundred base pairs (bp) in length, with average length varying from 340 bp to 720 bp across the six cell types. We extended the enhancers to be 3 kbp in length by including adjustable flanking regions, both for augmentation of positive samples and for more informative feature extraction with the use of the surrounding sequence context. The enhancers are fitted to a uniform length with the extensions, as sequences of fixed sizes are needed as input to our model. The original annotated promoters are mostly 1-2 kbp in length, with average varying from 450 bp to 1.96 kbp across cell types. Promoters are similarly fitted to a fixed window size of 2 kbp. If the original region is shorter than 3 kbp (for enhancer) or 2 kbp (for promoter), random shifting of the flanking regions is performed to get multiple samples. If it is longer than 3 kbp or 2 kbp, segments of the fixed length are randomly sampled from the original sequence.

For negative pairs, enhancers and promoters are also fixed to the window sizes of 3 kbp and 2 kbp, respectively, in the similar approach performed over the positive pairs, but without augmentation. The numbers of positive pairs, augmented positive pairs, and negative pairs on each cell line and the combined cell lines are listed in Table 1.

**Table 1:**
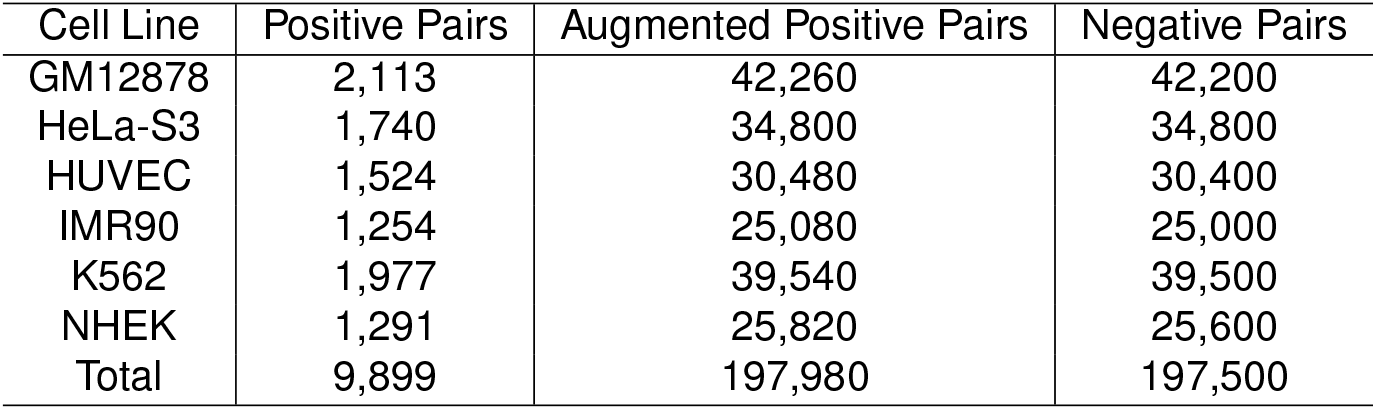
Number of positive sample, augmented positive sample, and negative sample counts, for each cell line.

### SPEID can effectively predict EPIs using sequence features only

We compared our prediction results to two state-of-the-art models using only information from individual cell type, TargetFinder [12] and PEP [15]. TargetFinder predicts EPI based on many functional genomic signals and annotations, including DNase-seq, DNA methylation, TF ChIP-seq, histone marks ChIP-seq, CAGE, and gene expression data. TargetFinder uses a dataset of enhancer and promoter pairs, labeled as interacting (positive) or non-interacting (negative). This dataset has 3 variants, which use features from different regions: Enhancer/Promoter (E/P) uses only annotations within the enhancer and promoter, Extended Enhancer/Promoter (EE/P) additionally uses annotations within an extended 3kbp flanking region around each enhancer, and Enhancer/Promoter/Window (E/P/W) additionally uses annotations in the region between the enhancer and promoter. PEP uses sequence-based features only and was trained and tested using the same collection of enhancers and promoters as TargetFinder (in all three variants). Note that in SPEID, our input sequences only include the surrounding sequences of the enhancer and promoter (as we discussed above). We did not compare SPEID with RIPPLE mainly because RIPPLE was trained on features similar to the EE/P dataset, but it is not compatible with the E/P/W data that we use here, and, furthermore, PEP was previously shown to consistently outperform RIPPLE in all cell lines on the EE/P dataset [15]. We therefore only directly compared with TargetFinder and PEP. SPEID and PEP both only consider sequence-based features, but SPEID has one advantage from a methodology stand-point. Since PEP performs separate feature extraction and prediction steps, the prediction model loses information about the contexts of features. In contrast, SPEID, which performs prediction directly from the sequences, can leverage the additional contextual information, such as relative positions of the features.

Fig. 2 shows the comparison of prediction performance between our SPEID method, the best PEP model, and the best TargetFinder model, on each of six different cell types, under each of the following performance metrics: (1) AUROC (area under receiver operating characteristic curve); (2) AUPR (area under precision-recall curve); and (3) *F*_1_ score (harmonic mean of precision and recall). AUROC and AUPR have the advantage that they do not depend on a particular classifier threshold. For *F*_1_, we used a classifier threshold that performed best on a predetermined validation data subset (10% of the training set, disjoint from the test set). We found that, although results vary across different cell lines, SPEID performs comparably to the most competitive variants of TargetFinder and PEP. Detailed numerical results and performance of other variants of TargetFinder and PEP are shown in Table S1. In summary, our results suggest that sequence contains important information that can determine EPI, and, if we are given the locations of enhancers and promoters for a particular cell type, our SPEID model can effectively predict EPI using sequence features only.

**Figure 2:**
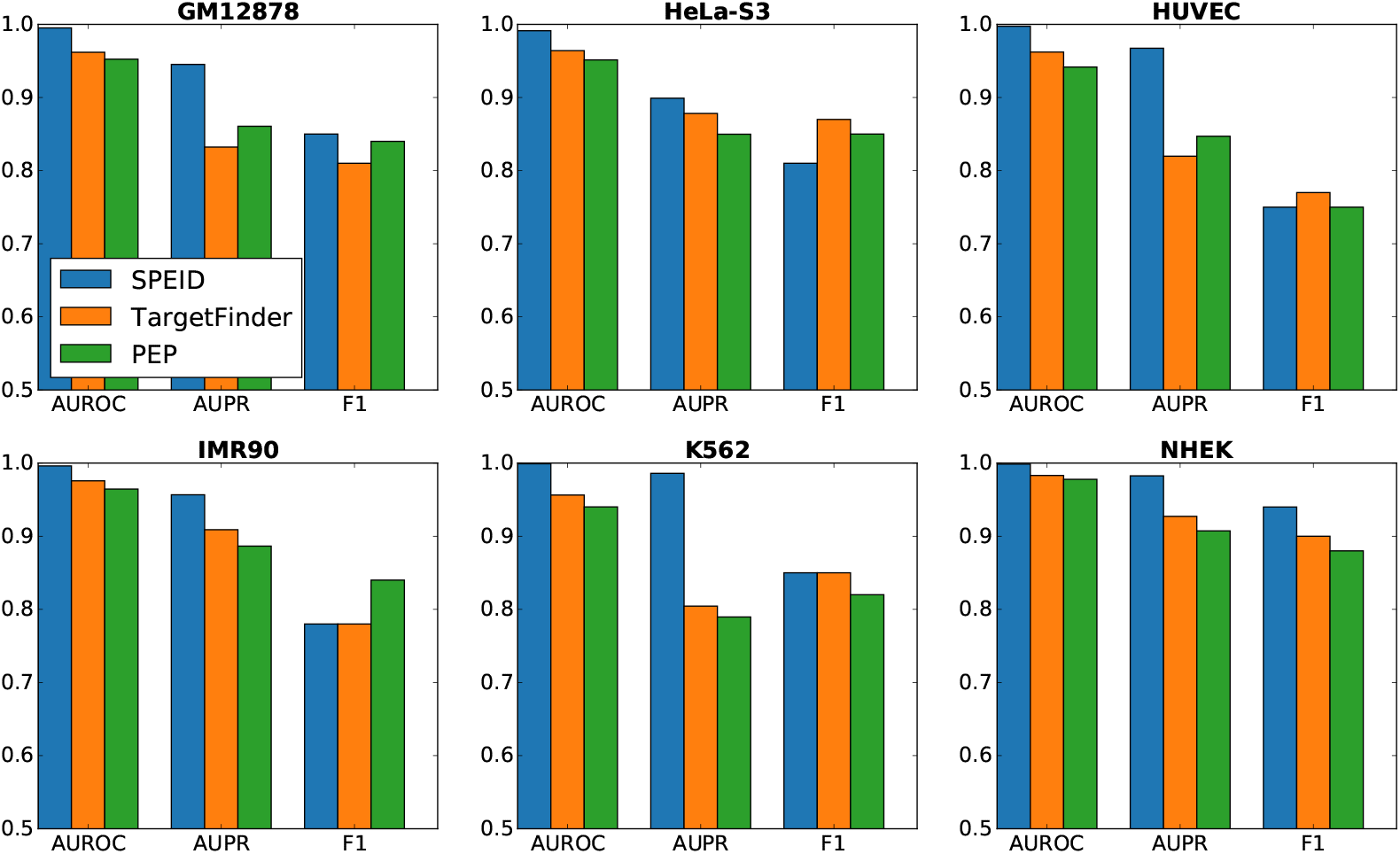
Prediction results of SPEID, TargetFinder’s E/P/W model, and PEP’s Integrated model in each cell line, as estimated by 10-fold cross-validation. AUROC, AUPR, and *F*_1_ are shown.

### Evaluating the importance of known sequence features using *in silico* mutagenesis

A major motivation for accurately predicting EPI from sequence is being able identify features of the DNA sequence that determine EPI. Unfortunately, unlike simpler models, deep learning models do not directly encode the features that they use to make their predictions. Nonlinearities in the deep network mean that ‘weights’ do not necessarily reflect ‘importance’, as in a linear regression model, and dropout regularization promotes a distributed representation of features within each layer of the network, such that small portions of learned features tend to be encoded in redundant fragments. Consequently, important features are difficult to extract directly from the network. Our main approach is therefore to study how changes in input sequences affect the predictions of SPEID for those sequences, a technique known as *in silico* mutagenesis. The mechanism underlying *in silico* mutagenesis is straightforward: for any particular pair of enhancer and promoter sequences, we can query SPEID with any mutation of those sequences to see whether this increases or decreases the predicted EPI probability. This allows us to predict how alterations to sequence affect EPI at nucleotide resolution.

While there are many ways this can be leveraged to identify important sequence features, we focused on measuring importance of known sequence motifs from the HOCOMOCO Human v10 database [30], which includes 640 motifs for 601 human TFs. Due to a high degree of redundancy/similarity of many of the motifs in HOCOMOCO, we first clustered the motifs (using the same approach described in Supplementary Methods A.2 of [15]), resulting in 503 motif clusters (including 427 single motifs and 76 small clusters of 24 motifs). For each motif cluster, in each of enhancers and promoters, we measured the change in prediction accuracy when all occurrences of motifs in that cluster in the test cross-validation fold were replaced with random noise (see Methods for more details). The average (over occurrences of that motif cluster) drop in prediction performance was then used as a measure of feature importance. Fig. S1 shows the distributions of estimated feature importance across all 503 motifs clusters, for each cell line, in each of enhancers and promoters. Since our measure of feature importance is an empirical average, the central limit theorem suggests that, if all features were equally important, all distributions in Fig. S1 would be approximately normal. However, most exhibit apparent positive skew, indicating the presence of a small number of highly positive feature importance values. In general, we found that enhancer features tended to be more important, consistent with the results in [12] and [15]. The top features according to SPEID’s feature importance scores also correlate significantly with those found by PEP [15], as shown in Table 2.

**Table 2:** Number of TF clusters (out of 503) predicted by both SPEID and PEP, only SPEID, and only PEP, to be in the top 10% feature importance in enhancers and promoters, in each cell line. Rows do not always sum to 50 due to exclusion of ties at the 10% cutoff, especially in PEP, whose feature importance scores are given in increments of 5%. For comparison, we have also provided chance values (e.g., if SPEID’s feature importance scores were randomly shuffled).

| Cell Line | Top 5% in both | Only in SPEID | Only in PEP |
| --- | --- | --- | --- |
| GM12878 | 22 | 28 | 26 |
| HeLa-S3 | 16 | 34 | 22 |
| HUVEC | 20 | 29 | 27 |
| IMR90 | 23 | 27 | 26 |
| K562 | 18 | 32 | 30 |
| NHEK | 14 | 36 | 17 |
| Chance | 5 | 45 | 45 |

To identify features that were consistently important across cell lines, we ranked features by importance within each cell line, and then averaged this rank across cell lines. Fig. 3 shows the 20 most important features (according to this average rank) in both enhancers and promoters, as well as their importance rank in each cell line. Table S2 shows the correlations between feature importance across different cell lines. The correlations are significantly positive, ranging from 0.24 to 0.37, suggesting that a considerable number of TF motifs play shared roles across multiple cell lines, while either Δ(*M*) is quite noisy or many motifs are important in some cell lines but not in others. The potentially important TFs include known ones such as CTCF, as well as a number of TFs whose roles in EPI have not been well studied at present, such as SRF, JUND, SPI1, SP1, EBF1, and JUN, which were also reported in [12]. Some of the highly predictive corresponding motif features discovered by SPEID are also consistent with evidence from existing studies. For example, SPEID ranks the motif of BCL11A in the top 5% important feature in both enhancer and promoter regions on average across different cell lines. Studies have shown that BCL11A could modulate chromosomal loop formation [31]. SPEID also ranks ZIC4, E2F3 and FOXK1 motifs as having top 5%, top 10%, and top 5% feature importance, respectively, in enhancer regions. ZIC4 and E2F3 are factors known to interact with enhancers [32], and their motifs are both found to be enriched in cohesin-occupied enhancers together with the CTCF motif [33]. FOXK1 has been shown to localize in both enhancers and promoters [34]. SPI1 (PU.1) is found by SPEID to be a corresponding motif feature with top 5% feature importance in promoter regions. Studies have revealed the important role of SPI1 in gene regulation [35, 36]. These results demonstrate the potential of SPEID in identifying important TF motif sequences involved in EPI without using any known TF motif information.

**Figure 3:**
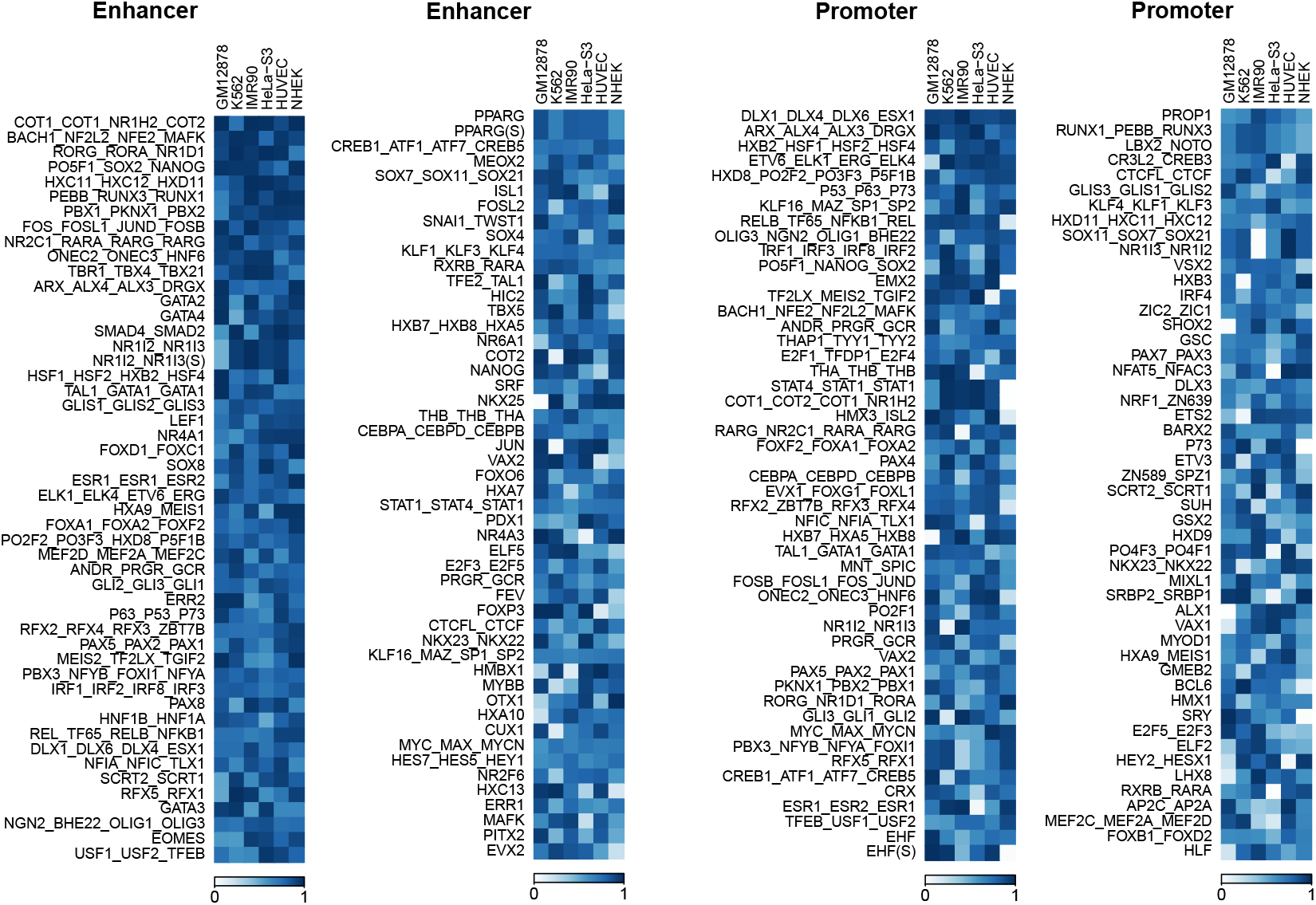
Feature importance in each cell line, for 100 features with highest average importance rank, sorted by average rank.

### Convolution features in SPEID reflect important TFs that mediate EPIs

Besides *in silico* mutagenesis, another useful way of understanding the features related to TF binding learned by SPEID is to compare the patterns of the convolutional kernels learned during training to known TF binding motifs (although SPEID likely also captures other informative features that do not match TF motifs). Using a similar procedure as in [20] and [19], we converted each kernel into a position frequency matrix (PFM). In short, this involves reconstructing the rectified output of the convolutional layer on each input sample sequence to identify subsequence alignments that best match each kernel, and then computing PFMs from these aligned subsequences (we used the same approach described in Supplementary Section 10.2 of [19]). We then used the motif comparison tool Tomtom 4.11.2 [37] to match these PFMs to known TF motifs from the HOCOMOCO Human v10 database.

Due to non-linearities in the deep network and the use of dropout during training, it is not obvious how to measure importance of specific convolutional features to prediction. Dropout encourages the model to develop redundant representations for important features, so that they are consistently available. As a result, we cannot measure importance of a convolutional kernel in terms of the change in prediction performance when holding that convolutional kernel out of the model directly. For the same reason, however, one measure of a feature’s importance is the redundancy of that feature’s representation in the model. Specifically, when using a dropout probability of 50%, the probability of a convolutional feature being available to the model is 1 − 2^−*r*^, where *r* is the number of copies of that convolutional feature, so that *r* is expected to be larger for more important features. As an example, the most frequently observed motif was the binding motif pattern of MAZ, which matched with 28 promoter convolution kernels. Note that MAZ was similarly reported as “of high importance” in promoter regions by TargetFinder. With this reasoning, we pruned the number of motif matches to report TFs that independently matched with at least 3 kernels in SPEID with an *E*-value of less than *E* < 0.5 according to Tomtom.

In Table 3, for each cell line, we give the number of motifs identified by SPEID using the approached introduced above, as well as the numbers of features found to be in the top 50% of importance by TargetFinder. Care must be taken when comparing the motifs discovered by SPEID with the features found important by TargetFinder; the latter used many features such as histone marks and gene expression data that lack corresponding motifs, and, even among TF features, many do not have corresponding motifs in the HOCOMOCO database. Furthermore, TargetFinder focused on TF ChIP-seq signals as features not only in the enhancer and promoter regions but also in the window region between them, while SPEID only uses sequence features from the input enhancer and promoter sequences and their flanking regions. Indeed, the importance of another feature, as measured by TargetFinder, is a function of the other features available to the model, and avoiding this subjectivity is an additional strength of SPEID. However, among the features that can be compared, the results suggest many commonalities between the findings of SPEID and TargetFinder. For example, of the 27 K562 enhancer motifs we discovered with corresponding features in TargetFinder, 23 were in the 30% of features considered most important by TargetFinder. Table 4 shows TFs with motifs discovered by SPEID, in two cell lines GM12878 and K562 (the two cell types with the largest number features in TargetFinder having corresponding motif, as well as the largest EPI datasets). These results further demonstrate the capability of SPEID in identifying important sequence features involved in EPI.

**Table 3:** Number of potentially important TFs in enhancers involved in EPIs identified by SPEID, TargetFinder, or both. Here we consider the top 50% most informative features from TargetFinder as important. The two methods have the largest overlap in GM12878 and K562 cell lines, likely because TargetFinder used many more TF ChIP-seq signals for these two cell lines.

| Cell Line | Predicted important in both | Only in SPEID | Only in TargetFinder |
| --- | --- | --- | --- |
| GM12878 | 22 | 9 | 53 |
| HeLa-S3 | 13 | 15 | 37 |
| HUVEC | 1 | 14 | 7 |
| IMR90 | 4 | 31 | 16 |
| K562 | 27 | 26 | 85 |
| NHEK | 0 | 16 | 5 |

**Table 4:**
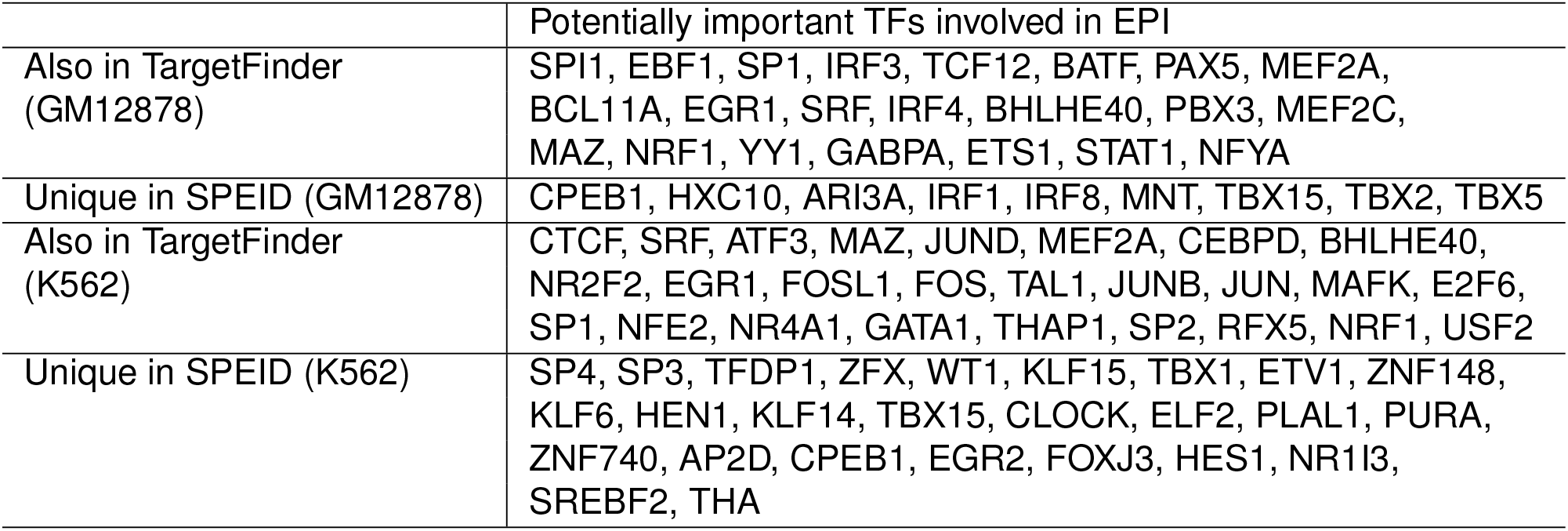
Predicted potentially important TFs in enhancers involved in EPIs from SPEID in GM12878 and K562.

|  |  |
| --- | --- |
|  | Potentially important TFs involved in EPI |
| Also in TargetFinder (GM12878) | SPI1, EBF1, SP1, IRF3, TCF12, BATF, PAX5, MEF2A, BCL11A, EGR1, SRF, IRF4, BHLHE40, PBX3, MEF2C, MAZ, NRF1, YY1, GABPA, ETS1, STAT1, NFYA |
| Unique in SPEID (GM12878) | CPEB1, HXC10, ARI3A, IRF1, IRF8, MNT, TBX15, TBX2, TBX5 |
| Also in TargetFinder (K562) | CTCF, SRF, ATF3, MAZ, JUND, MEF2A, CEBPD, BHLHE40, NR2F2, EGR1, FOSL1, FOS, TAL1, JUNB, JUN, MAFK, E2F6, SP1, NFE2, NR4A1, GATA1, THAP1, SP2, RFX5, NRF1, USF2 |
| Unique in SPEID (K562) | SP4, SP3, TFDP1, ZFX, WT1, KLF15, TBX1, ETV1, ZNF148, KLF6, HEN1, KLF14, TBX15, CLOCK, ELF2, PLAL1, PURA, ZNF740, AP2D, CPEB1, EGR2, FOXJ3, HES1, NR1I3, SREBF2, THA |

### Predicting effects of somatic mutations on EPI in melanoma

To further demonstrate the usefulness of EPI prediction, we applied SPEID to study the effects of somatic mutations on EPI in melanoma patients. We used a somatic mutation dataset involving 183 melanoma patients [25, 38]. The positions and types of DNA mutations on the whole genome were identified for each of the patients. We extracted DNA sequences from the same pairs of interacting enhancer and promoter regions as we used in the NHEK (Normal Human Epidermal Keratinocytes), which serves as the normal skin cell here. For each of 183 patients and each of the 1,291 positive EPI loops in the NHEK training dataset, we used SPEID to predict the interaction likelihood of that EPI given the patient’s mutated enhancer and promoter sequences. For certain EPI pairs, SPEID predicted a significantly lower interaction likelihood (caused by somatic mutations) than for the original sequences, suggesting that those EPI might be potentially reduced in the patient sample. In particular, we identified those EPIs that are predicted as being potentially reduced in multiple patients. By using a threshold to select loops with more significant decrease of interaction likelihood predicted by SPEID, we identified 178 EPIs that are reduced in at least one of the 183 patients. We identified 61 EPIs that are likely to be reduced in at least two patients and 27 EPIs which are likely to be reduced in at least three patients (see Supplementary Table S3 for the list).

We then investigated whether the mutations might interfere with TF motifs that SPEID had previously identified as having high importance in NHEK. We first identified TF motifs in the normal sequences of the 1,291 positive EPIs, using the results of motif scanning we performed in the *in silico* mutagenesis. For each possibly reduced EPI of each patient, we searched for the motifs that are overlapping with at least one somatic mutation in the patient sample. We ranked the motifs by the total number of mutations they encountered across different enhancer/promoter pairs of different patients. We observed that the motifs of MAZ, SP1, and EGR1 are the top 3 TF motifs with the highest frequent mutations. MAZ, SP1, and EGR1 are also predicted by SPEID to be of high feature importance in NHEK. We then examined how the mutations may affect the likelihood of a TF binding site, using the Position Weight Matrices (PWM) of motifs from the HOCOMOCO human v10 database. For each mutated position, we compared the position weight of the original nucleotide and the nucleotide after mutation. We observed that in the predicted reduced EPIs, 86% of the mutations within MAZ motifs, 78% of the mutations within SP1 motifs, and 63% of the mutations within EGR1 motifs have induced decrease of the position weight, respectively. For example, in Fig. 4 we show that the EPI connecting the enhancer at chr19:6,516,000-6,516,200 and the promoter at chr19:6,737,600-6,737,800 in NHEK is predicted by SPEID to be reduced in 5 patients. There are 5 mutations within the enhancer or promoter regions of this EPI of 5 patients, of which 3 overlap with the motifs of SP1 or NFKB1. In Fig. 4 we show one somatic mutation in Patient # DO220903 where the estimated reduction likelihood ranked in the top 0.2% of all the EPIs in NHEK. This analysis provides a proof-of-principle to demonstrate the potential of applying SPEID to identify somatic non-coding mutations that may reduce or disrupt important EPIs.

**Figure 4:**
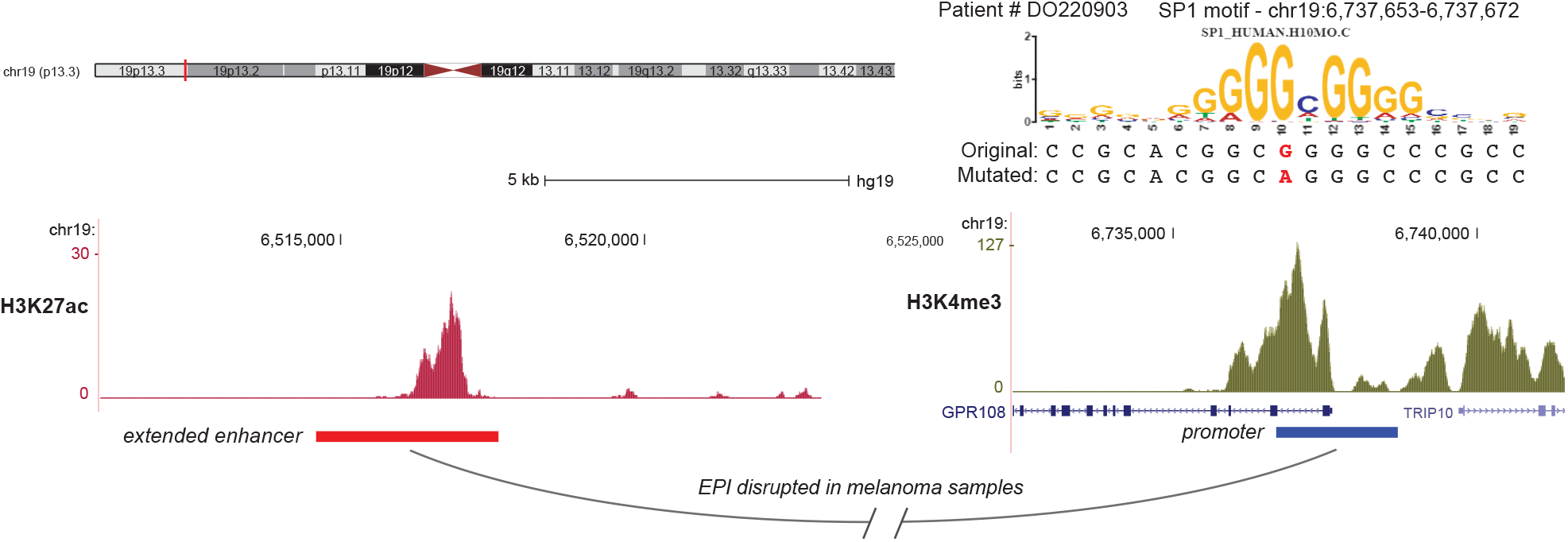
Example of a possibly reduced EPI (extended enhancer at chr19:6514600-6517600, promoter at chr19:6736700-6738700) with mutation occurring within the motif of SP1 in the promoter region shown. The enhancer region is extended to be 3kb in length and the promoter region is 2kb. The estimated reduction likelihood of this EPI with the mutation is ranking top 0.2% of all the positive EPIs in the NHEK cell line.

## DISCUSSION

Long-range interaction between enhancers and promoters is one of the most intriguing phenomena in gene regulation. Although new high-throughput experimental approaches have provided us with tools to identify potential EPIs genome-wide, it is largely less clear whether there are sequence-level instructions already encoded in our genome that help determine EPIs. In this work, we have developed, to the best of our knowledge, the first deep learning model, SPEID, to directly tackle this question. The resulting contributions are as follows: (1) We have shown that sequenced-based features alone can indeed effectively predict EPIs, given the genomic locations of putative enhancers and promoters in a particular cell type; SPEID achieves performance competitive with the state-of-the-art method TargetFinder that uses a large number of functional genomic signals instead of sequence features. (2) By learning important sequence features in a supervised manner, deep models can outperform methods such as PEP that use either manual or unsupervised feature extraction (i.e., independent of the classification task). (3) The deep learning framework in SPEID provides a useful predictive model for studying genomic sequence-level interactions by extracting relevant sequence information, representing an important conceptual expansion of the application of deep learning models in regulatory genomics. (4) Despite the complexity of deep learning models, important sequence features can be extracted, both by inspection of the convolutional kernels, and via *in silico* mutagenesis. (5) The predictions from deep learning models can be applied downstream to investigate connections between non-coding mutations and diseases.

However, methods used with PEP [15] and TargetFinder [12] and our *in silico* mutagenesis method used with SPEID have two main differences:

1. When measuring feature importance, TargetFinder and PEP are restricted to hand-picked features, rather than arbitrary sequence features. ^1^ This is important because (a), unlike our methods, these cannot be used for measuring importance of novel features, and (b), perhaps more importantly, *the importance of a feature in a predictive model depends on the other features available to the model*. As a result, our importance measure, based on using the entire sequence rather than hand-picked features, is more objectively interpretable than those of PEP and TargetFinder. Specifically, it measures the importance of sequence features *relative to the rest of the sequence*, rather than relative to the other hand-picked features available to the model.
2. While all these approaches rely on heuristics to search the huge space of possible features combinations, the importance measures used with PEP and TargetFinder are “additive” (i.e., they measure benefit of adding a feature), while our measure is “subtractive” (i.e., it measures cost of removing a feature). Said another way, PEP and TargetFinder identify features that are *sufficient* for prediction, whereas SPEID identifies features that are *necessary* for prediction, given the rest of the sequence. These approaches are complementary.

Both these differences make our feature importance measure more conservative than the measure of PEP; although the measures are strongly correlated, in Fig. S2, most points at which PEP and SPEID differ lie above the diagonal.

There are a number of directions in which our method can be further improved. First, our current ability in determining the potentially informative features remains limited. Although we were able to identify some informative TFs that may play roles in mediating EPIs in a certain cell type and that were also identified from TargetFinder, a significant proportion of sequence features from our model cannot be easily interpreted and their contributions are also hard to evaluate. One approach may be to apply very recently developed methods such as DeepLIFT [39] and deep feature selection [40] for measuring importance of and selecting among different convolutional features, to identify those that might mediate EPI within or across cell lines.

However, *in silico* mutagenesis is a versatile tool, with several potential applications beyond just measuring predictive importance of given sequence features. For example, it can also provide more information about the role of important sequence features. A reduction in prediction accuracy associated with removing a particular feature might be driven by increasing the false positive rate or by increasing the false negative rate. The former would suggest that the presence of the feature promotes interactions, whereas the latter would suggest that it represses interactions. However, since *in silico* mutagenesis is also computationally intensive (in each cell line and each of enhancers and promoters, a total of around 400 hours are needed to complete). Thus, while it may have other applications in this context, we have restricted ourselves to measuring feature importance. ^2^

The current SPEID framework depends on sequence features that are cell-type specific, and is unable to automatically capture relevant sequence features operating across cell types. As more positive EPI samples and data in additional cell types become available, more work is needed to determine exactly what, if any, sequence features mediate EPI consistently across cell types. Finally, though we demonstrated that sequence features alone can effectively predict EPI, it would be important to explore the optimal combination of sequence-based features and features from functional genomic signals to achieve the strongest predictive power in a cell-type specific manner. Such an approach would be useful in understanding the genetic and epigenetic mechanisms that determine EPIs, and their variation across different cell types. Our work lays foundation for this by providing a new framework to potentially decode important sequence determinants for long-range gene regulation.

## MATERIALS AND METHODS

### Model framework of SPEID

As shown in Fig. 1, the main layers of the network are pairs of layers for convolution, activation, and max-pool layers, respectively, together with a single recurrent long short-term memory (LSTM) layer, and a dense layer.

#### Input, convolution, and max-pooling

The first layers of the network are responsible for learning informative subsequence features of the inputs. Because informative subsequence features may differ between enhancers and promoters, we train separate branches for each. These features might include, for example, TF protein binding motifs and other sequence-based signals. Each branch consists of a *convolution layer* and a rectified linear unit (ReLU) *activation layer* [41], which together extract subsequence features from the input, and a *pooling layer*, which reduces dimensionality. Recall that each sequence input is a 4 × 3000 matrix (for enhancer) or 4 × 2000 matrix (for promoter), with a one-hot encoding (e.g., ‘A’ is (1,0,0, 0)^*T*^, ‘G’ is (0,1, 0, 0)^*T*^, ‘C is (0, 0,1,0)^*T*^, and ‘T’ is (0, 0, 0,1)^*T*^). The convolution layer consists of an array of 200 ‘kernels’, 4 × 40 signed weight matrices that are convolved with the input sequence to output a sequence of ‘scores’, indicating how well the kernel matches with each 40bp window of the input sequence at each possible offset. More precisely, each kernel is a matrix *K* ∈ ℝ^4×40^, and, for each one-hot encoded input matrix *X* ∈ {0, 1}^4×*L*^ and each offset *i* ∈ {0,…, *L —* 40} (where *L* is the length of the input sequence), the convolution layer outputs the matrix inner product:

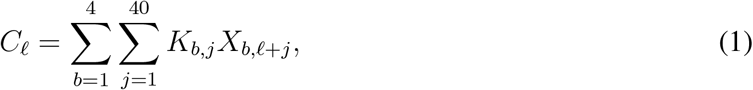

between *K* and the 40bp submatrix of *X* at offset *l*. A higher (more positive) *C_l_* indicates a better match between *K* and the *l^th^* offset subsequence of the input.

*C_l_* is then passed through a ReLU activation *R*(*x*) = max{0, *x*}, which propagates positive outputs (i.e., sequence ‘matches’) from the convolution layer, while eliminating negative outputs (i.e., ‘non-matches’). Of the various non-linearities that can be used in deep learning models, ReLU activations are the most popular, due to their computational efficiency, and because they naturally sparsify the output of the convolution layer to only include positive matches [42].

Max-pooling then reduces the output of the convolution/activation layer by propagating only the largest output of each kernel within each ‘stride’ (i.e., 20 bp window), effectively outputting the ‘best’ alignment of each kernel within each stride. Specifically, it reduces the sequence *R*(*C*_1_), *R*(*C*_2_),…, *R*(*C*_L–40_) of length *L* — 40 to the subsampled sequence:

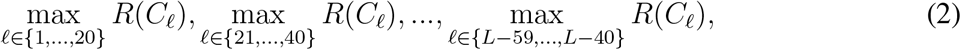

of length (*L* — 40)/20. Pooling is especially important in EPI prediction because of the long input sequence (5kbp, as compared to 1kbp used when predicting function of single sequence variants [17, 20]). Here we use the following parameters: Number of Kernels: 200, Filter Length: 40, *L_2_* penalty weight: 10^−5^, Pool Length: 20, Stride: 20.

Before feeding into the next layer, the enhancer and promoter branches are concatenated into a single output. The remaining layers of the network act jointly on this concatenation, rather than as disjoint pairs of layers, as in the previous layers (see Fig. 1).

#### LSTM, dense layer, and final output

The next layer is a *recurrent long short-term memory (LSTM) layer* [43], responsible for identifying informative combinations of the extracted subsequence features, across the extent of the sequence (for the internal mechanism of an LSTM, see [44] for the detailed explanation of the particular LSTM implementation we use). As a brief intuition, the LSTM outputs a low-dimensional weighted linear projection of the input, and, as the LSTM sweeps across each element of the input sequence, it chooses, based on previous inputs, the current input, and weights learned by the model, to add or exclude each feature in this lower dimensional representation. This layer is bidirectional; that is it sweeps from both left to right and right to left, and the outputs of each direction are concatenated for a total output dimension of 100.

The final *dense layer* is simply an array of 800 hidden units with nonlinear (ReLU) activations feeding into a single sigmoid (i.e., logistic regression) unit that predicts the probability of an interaction. That is, if *y* ∈ ℝ^100^ denotes the output of the LSTM layer, then then final output is a predicted interaction probability *S*(*υ*^*T*^ *R*(*Wy*)) ∈ (0, 1), where *W* ∈ ℝ^800×100^ and *υ* ∈ ℝ^100^ are learned weights, *R* denotes ReLU activation (applied to each component of *Wy*), and 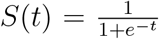 denotes the sigmoid function.

Similar (albeit simpler) architectures have been used for the related problem of predicting function of non-coding sequence variants [17, 20]. In fact, our use of a recurrent LSTM layer rather than a hierarchy of convolutional/max-pooling layers is inspired by the architecture of the DanQ model [20], which suggests that the LSTM is better able to model a “regulatory grammar” by incorporating long range dependencies between subsequences identified by the convolution layer. However, our method solves a fundamentally different problem – predicting interactions between sequences rather than predicting annotations from a single sequence. Hence, our model has a branched architecture, taking two inputs and producing a single classification, rather than a sequential architecture. Because the data for this problem are far sparser, we require a more careful training procedure, as detailed in the next section. There are also several finer distinctions between the models, such as our use of batch normalization to accelerate training and weight regularization to improve generalization.

#### Other model and implementation details

We implemented our deep learning model using Keras 1.1.0 [45]. The model was trained in mini-batches of 100 samples by back-propagation, using binary cross-entropy loss, minimized by Adam [46] with a learning rate of 10^−5^. The pre-training and re-training phases lasted 32 epochs and 80 epochs, respectively. The training time was linear in the sample size for each cell line, taking, for example, 11 and 6 hours for pre-training and retraining phases, respectively, on K562 data, on an NVIDIA GTX 1080 GPU.

Due to SPEID’s many hyperparameters and the computational overhead of training the model, we tuned hyperparameters using the full data set from only one cell line (K562), and then used the same hyperparameter values for all other cell lines. For this reason, we emphasize our results on the remaining 5 cell lines, where the trained model is entirely independent of the test data. Here, we list the full range of model parameters we tried, with the finally selected values in **bold**:

1. Convolution kernel lengths: 26, **40**, 50
2. Number of convolution kernels: 100, **200**, 320, 512, 1024
3. Number of neurons in dense layer: 600, **800**, 1000
4. LSTM output dimension: 50, **100**, 200, 500
5. Dropout probability: 0.25, **0.5**, 0.6
6. L_2_ regularization weight: 0, 10^−6^, **10**^−3^

### Addressing potential overfitting

When training very large models such as deep networks, potential overfitting is a concern. This is particularly relevant because our datasets are small (< 10^4^ positive samples per cell line) compared to the massive data sets often used to train deep networks (e.g., with imaging or text data). We employ multiple approaches to prevent or mitigate overfitting, both within the deep learning model, and in our training and evaluation procedures.

Firstly, note that we provide results on six independent data sets from different cell lines. While we experimented with multiple deep network models, utilizing different layers and training hyperparameters, we did this based on only test results from K562. In particular, results on the remaining 5 cell lines are independent of model selection. Second, by randomly shifting positive inputs, our data augmentation (see below) specifically prevents the model from overfitting to any region of the input. Finally, our deep network model itself incorporates three tools to reduce overfitting during training: batch normalization, dropout, and *L*_2_ regularization.

Batch normalization [47] is the process of linearly normalizing the outputs of neurons on each training batch to have sample mean 0 and standard deviation 1. That is, if the *i*-th batch consists of 100 samples for which a particular neuron gives outputs *N*_*i*,1_, …, *N*_*i*,100_, then we replace these outputs with normalized outputs

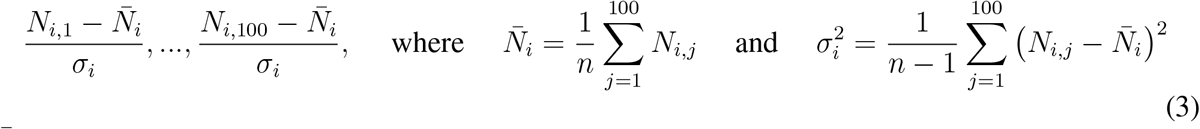

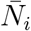 and 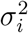 are the sample mean and sample variance, respectively. Batch normalization combats overfitting by limiting the range of non-linearities (in our case, ReLU function) in the network, and also accelerates training (i.e., reduces the number of epochs till convergence) by restricting the input space of downstream neurons. We batch-normalize the outputs of 4 layers in the network: the max-pooling layer, the LSTM layer, the dense layer (i.e., before the ReLU activation), and the ReLU activation (i.e., before the final sigmoid classifier). Note that, during prediction, batch means and variances are replaced with population means and variances, which are computed while training over all batches.

Dropout, a common regularization technique in neural networks, refers to randomly ‘dropping’ (i.e., setting to zero) the output of a neuron with some fixed probability *p*. That is, for each sample *i* and each neuron *j*, we sample a Bernoulli(p) random variable *D_ij_* and replace the output *N_ij_* with (1 — *D_ij_*)*N_ij_*. Applying dropout to a layer L_i_ prevents the subsequent layer L_i+1_ from overfitting to any subset of neurons Li in intermediate layers, and there by promotes a distributed representation, which can be thought of as *model averaging* (i.e., learning the average of many models, as in ensemble techniques). We apply dropout with *p* = 0.5 to the outputs of 3 layers: the max-pooling layer, the LSTM, and the dense layer (dropout is always applied after batch normalization). Note that dropout is only applied in training; all neurons’ outputs are used during testing.

Finally, we apply an *L*_2_-norm penalty to the kernels in the convolution layer and on the matrix of weights of the dense layer. As with the *L*_2_ penalty commonly applied in linear regression, this helps ensure that no particular weight becomes too large.

### Training procedure

Recall that our data set is highly imbalanced – there are many more negative (non-interacting) pairs than positive (interacting) pairs. In each cell line, there are typically 20 times more negative samples than positive samples. To combat the difficulty of learning highly imbalanced classes, we utilize a two-stage training procedure that involves pre-training on a data set balanced with data augmentation, followed by training on the original data.

#### Pre-training with data augmentation

Data augmentation is commonly used as an alternative to re-weighting data when training deep learning models on highly imbalanced classes. For example, image data is often augmented with random translations, scalings, and rotations of the original data [48]. In our case, because enhancers and promoters are typically smaller than the fixed window size we use as input, the labels are invariant to small shifts of the input sequence, as long as the enhancer or promoter remains within this window. By randomly shifting each positive promoter and enhancer within this window, we generated “new” positive samples. We did this 20 times with each positive sample, resulting in balanced positive and negative classes. In addition to balancing class sizes, this data augmentation has the additional benefit of promoting translation invariance in our model, preventing it from overfitting to any particular region of the input sequence.

#### Imbalanced training

Data augmentation results in a consistent training procedure for the network, allowing the convolutional layers to identify informative subsequence features and the recurrent layer to identify long-range dependencies between these features. However, in typical applications of predicting interactions, classes are, as in our original data, highly imbalanced. In these contexts, naively using the network trained on augmented data results in a very high false positive rate. Fortunately, this has relatively little to do with the convolutional and recurrent layers of the network, which correctly learn features that distinguish positive and negative samples, and this issue is largely due to the dense layer, which performs prediction based on these features. Hence, to correct for this, we only retrain the dense layer. We do this by “freezing” the lower layers of the network (i.e., setting the learning rate to 0), and then continuing to train the network as usual on the subset of the original imbalanced data that was used to generate the augmented data.

#### Summary of training procedure

The following procedure is repeated independently for each of the five cell lines we used for evaluation:

1. Begin with an imbalanced data set *A*.
2. Split *A* uniformly at random into a training set *B* (90% of *A*) and a test set *C* (10% of *A*).
3. Augment positive samples in *B* to produce a balanced data set *D*.
4. Train the model on *D*, using a small (10%) subset for model validation.
5. Freeze the convolution and recurrent layers of the model.
6. Continue training the dense layer of the model on *B*.
7. Evaluate on *C*.

We performed 10-fold cross-validation, repeat steps 2 through 7 for each of 10 disjoint training-test splits to reduce evaluation variance.

### Measuring feature importance with *in silico* mutagenesis

Here, we describe our procedure for using SPEID, together with *in silico* mutagenesis, to measure the importance of motifs in the HOCOMOCO database [30] as predictive features for EPI. For each motif *M* in the HOCOMOCO database, each cell line, and each of enhancers and promoters, we first used FIMO [49] to scan for all occurrences of *M* in our test set *D*. Next, we replaced each occurrence of *M* with random noise, in a copy *D’* of *D*. We then measured the prediction performance *P’(M)* of SPEID on *D’*, subtracted this from the performance *P* on *D*, and normalized by dividing by the number of occurrences *N(M)* of *M*. Our measure of the importance of motif *M* was then represented as:

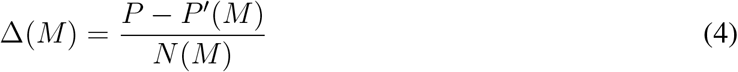

which measures, on average, how necessary occurrences of motif *M* are for predicting EPI. We used AUPR to avoid dependence on the classification threshold. To prevent biasing towards longer motifs (whose modification would change more of the input sequence), rather than mutating the exact sequence match identified by FIMO, we mutated a 20 bp window centered at the match center (as nearly all HOCOMOCO motifs are less than 20 bp long). If a match center was within 10 bp of the end of the input sequence, we mutated the 20 bp at that end of the sequence. For each motif, this procedure was performed separately for both enhancer and promoter inputs, producing two independent scores for each motif. To minimize variance in estimating Δ(*M*), we averaged estimates over each of 10 cross-validation folds, so that each occurrence of each motif in the dataset was mutated exactly once (since each sample occurs in the test set of exactly one CV fold).

## Acknowledgments

We thank the members of the Ma lab, especially Yang Zhang, Yuchuan Wang, Ruochi Zhang, and Dechao Tian, for helpful discussions. This work was supported in part by the National Science Foundation [1252522 to S.S., 1054309 and 1262575 to J.M.] and the National Institutes of Health [HG007352 and DK107965 to J.M.].

## Footnotes

^1^Although the PEP-Word module allows prediction of EPI from sequence without manual feature selection, [15] used only the PEP-Motif module, based on known motifs from HOCOMOCO, to measure feature importance.

^2^This procedure takes order O(nMC) time, where n is the average sample size per cell line, C is the number of cell lines, and M is the number of motifs. However, it also parallelizes well when multiple GPUs are available.

